# Hantaan virus replication is promoted via AKT activated mitochondria OXPHOS

**DOI:** 10.1101/2022.01.05.475173

**Authors:** Yuhang Dong, Xiaoxiao Zhang, Mengyang Li, Qikang Ying, Yunan Feng, Zhoupeng Li, Xingan Wu, Fang Wang

## Abstract

Oxidative phosphorylation (OXPHOS) is a vital pathway provides ATP for intracellular activities. Here, we found that Hantaan virus (HTNV) exploited mitochondria OXPHOS to assist its replication in host cells and Protein Kinase B/AKT played a major function in this process. Inhibiting AKT activation by BEZ treatment can inhibit HTNV replication and prevent the increase of OXPHOS level caused by HTNV infection. We also found that HTNV infection can promote AKT translocation to mitochondria, where AKT phosphorylates Polynucleotide phosphorylase (PNPT). Taken together, our research demonstrates that HTNV replication exploits OXPHOS in host cells and it increases OXPHOS function by AKT-PNPT interaction in mitochondria.

**IMPORTANCE:** Virus depends on metabolic pathways in host cells to favor its replication. This is a vital process which needs complicated host-virus interaction and targeting this process is a new strategy for antiviral drug development. Hantaan virus (HTNV) is the major pathogen which causes Hemorrhagic Fever with Renal Syndrome (HFRS) in China. However, there are neither effective therapeutic drugs nor FDA-licensed vaccine against HFRS, a deeper understanding of HTNV infection characteristics is of great significance for global public health and safety. This research means to elucidate the major metabolic pathway exploited by HTNV during its replication in host cells and its underlying molecular mechanism, which can enrich our understanding about HTNV biological characteristics and pathogenesis, also provide a new view on anti-HTNV drug development.

---

Hantaan virus (HTNV) is an enveloped virus which contains a tripartite negative-sense single-stranded RNA (ssRNA) genome, including small (S), medium (M), and large (L) genes, which encode nucleocapsid protein (NP), glycoprotein (GP) and viral RNA-dependent polymerase protein (RdRp) respectively. It belongs to the family Bunyaviridae and genus Hantavirus, and is the major pathogen of Hemorrhagic Fever with Renal Syndrome (HFRS) in China^[1] [2]^. HTNV replicates slowly in host cells and does not induce cytopathic effect (CPE), such characteristics may due to metabolic changes in host cells during its infection, which is still elusive by now.

Virus replication makes use of energy and metabolic materials from host cells^[3]^. Eukaryotes produce energy by glycolysis and oxidative phosphorylation (OXPHOS). After entry into host cells, viral protein can interact with key enzymes or signal molecules to mediate cell metabolic pathways to assist its replication.

Most viruses, like Hepatitis C (HCV)^[4]^, Adenovirus (Adv)^[3]^, Dengue virus (DENV)^[5]^, increase cellular aerobic glycolysis (Warburg Effect) to support its replication. The intermediate products from Warburg effect can be used to viral nucleic acid or amino acid synthesization directly. Some viruses, such as Human Cytomegalovirus (HCMV)^[6, 7]^, Rubella Virus (RV)^[8]^, Measles Virus (MV)^[9]^, exploit OXPHOS as a major pathway to provide energy for its replication. For example,, HCMV encoded non-coding RNA β2.7 can combine with mitochondria RC complex I directly, increasing complex I activity and ATP production^[10]^; HCMV infection can induce nuclear DNA-encoded PGC-1α^[6]^, NRF2^[11]^ expressions, which can promote mitochondria DNA (mtDNA) transcription.

Mitochondria is an independent, double-membrane organelle, it is critical for energy generation and is also recognized as gatekeepers in the control of cell survival, oxidative stress and innate immune^[12]^. mtDNA is a small circular genome, which encodes 2 rRNAs, 22 tRNAs, and 13 mRNA^[13]^.

AKT is an important kinase which mediates cell growth, proliferation and metabolism. A majority of viruses infection can activate AKT signal pathway to inhibit apoptosis or facilitate cell metabolic reprogramming^[20]^. AKT has 2 important phosphorylation sites: Thr^308^ which can be phosphorylated by phosphatidylinositol 3 kinase (PI3K)-dependent Kinase 1 (PDK1) ^[21]^, and Ser^473^ which can be phosphorylated by mammalian Target Of Rapamycin Complex 2 (mTORC2)^[22]^. PH domain of AKT has high affinity with phosphate inositol, which allows AKT access to any cellular membrane structure^[23]^. Whether AKT mediate mitochondria OXPHOS level positively^[24][25, 26]^ or negatively^[27, 28]^, which is dependent on different AKT-phosphorylated substrate in mitochondria under different situations. Our study means to delineate cellular metabolic change after HTNV infection and its underlying mechanism.

## Results

### HTNV infection increases OXPHOS of host cells Three

HTNV-permissive cells with different metabolic backgrounds, Human Umbilical Vein Endothelial cell (HUVEC) as non-tumorogenic cell line, mouse bone marrow derived macrophage (BMDM) as untransformed primary cells and human lung carcinoma cell line A549, were used for a comprehensive analysis of the metabolic changes during HTNV infection. Multiplicity of Infection (MOI) of 3 was chosen to achieve a high initial infection rate. All these cells are susceptible to HTNV and can reach a high replication level at 2 days post infection (dpi.) (Fig. S1).

Using Seahorse XFe24 Bioanalyzer, we examined OXPHOS level changes following HTNV infection, which was measured by oxygen consumption rate (OCR) (Fig 1). Significantly higher levels of basal and maximal respiration, spare respiratory capacity and ATP production were observed in HTNV-infected cells compared with mock-infected cells. Those increased indexes reflected a total OXPHOS level increase per cell, which was caused by enhancement of single mitochondria OXPHOS capacity or mitochondria quantity increase. During oxidation in mitochondrial respiratory chain, leaked proton can react with oxygen in mitochondria matrix, forming reactive oxygen species (ROS). mtROS is an index used to evaluate mitochondrial oxidation efficiency. Compared with the higher oxygen consumption after HTNV infection, little changed proton leak levels implied relatively higher proton pump efficiency, which also means increase of mitochondria oxidation function. We also detected mitochondria ROS (mtROS) by flow cytometry using a MitoSOX^™^ Red dye, no significant changes were found after HTNV infection (Fig. S2). Therefore, mitochondrial respiration enhancement after HTNV infection was caused by enhanced single mitochondria OXPHOS capacity rather than increased mitochondria quantity.

**Fig 1.**
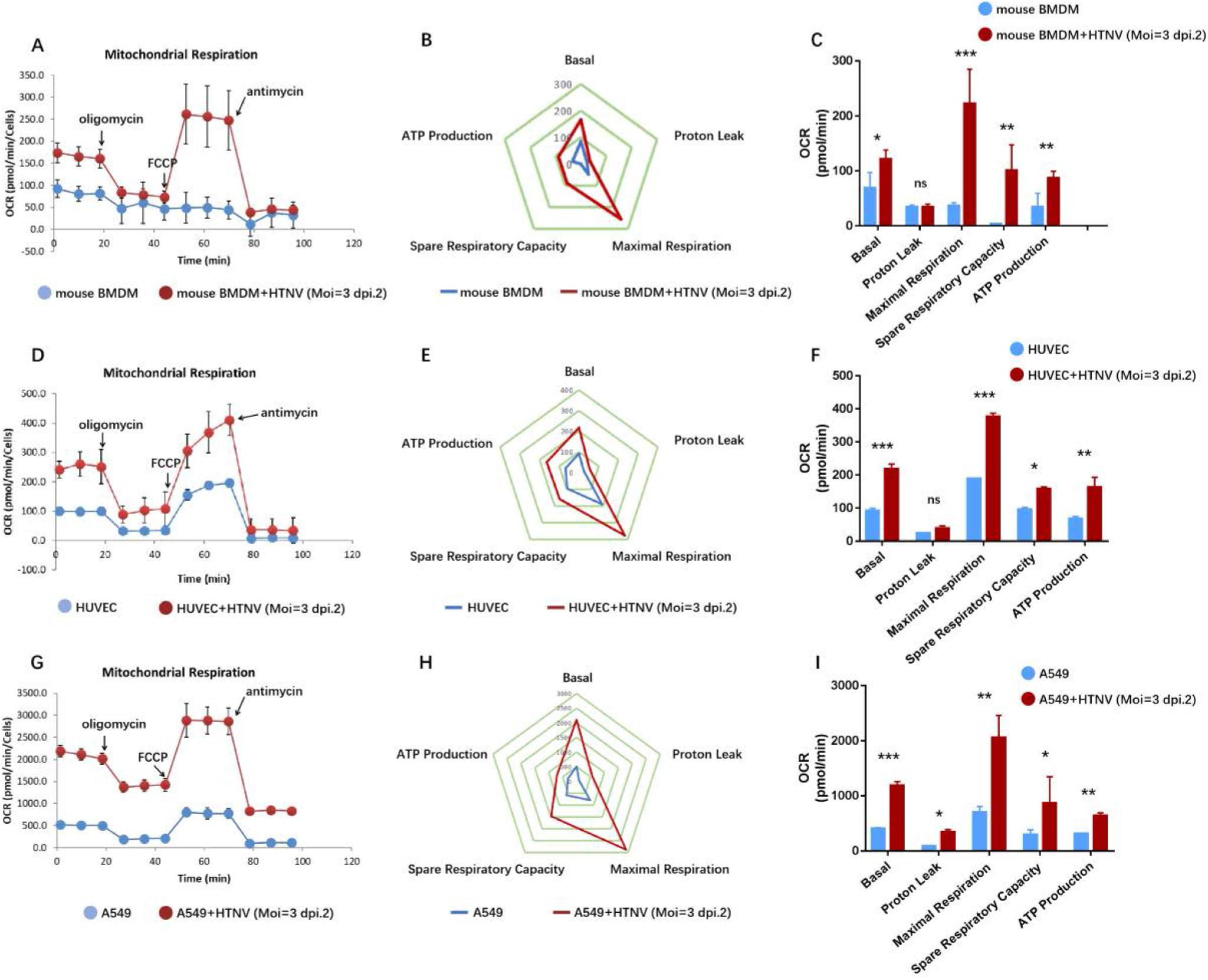
Mitochondria OXPHOS function increased obviously after HTNV infection. Mitochondria respiration was measured 2 days after HTNV infection using the Seahorse XFe24 Mito Stress Test (A: mouse BMDM; D: HUVEC; G: A549 cells). OCR was detected every 8.5 minutes and the dynamic change was presented as line chart. All OCR data were normalized and the data were from at least three independent experiments. The results were also analyzed and expressed via radar map (B, E, H) and column diagram (C, F, I). Error bars indicate the standard errors of the mean (SEM) of at least triplicates. (Student’s t test, *, P< 0.01; **, P<0.001; ***, P<0.0001; ns: no significance.)

We next tested whether inhibitors of OXPHOS or glycolysis could influence HTNV replication. We treated A549 cells with Rotenone (mitochondria electron transport chain inhibitor) or 2-DG (a glucose analog that blocks early glycolysis). We found that after Rotenone treatment, HTNV replication was decreased obviously (Figure 2A), but 2-DG treatment had a relatively minor effect on HTNV replication (Figure 2B). Thus, we concluded that HTNV replication can be inhibited once mitochondria OXPHOS was inhibited, which means that HTNV replication depended on OXPHOS.

**Fig 2.**
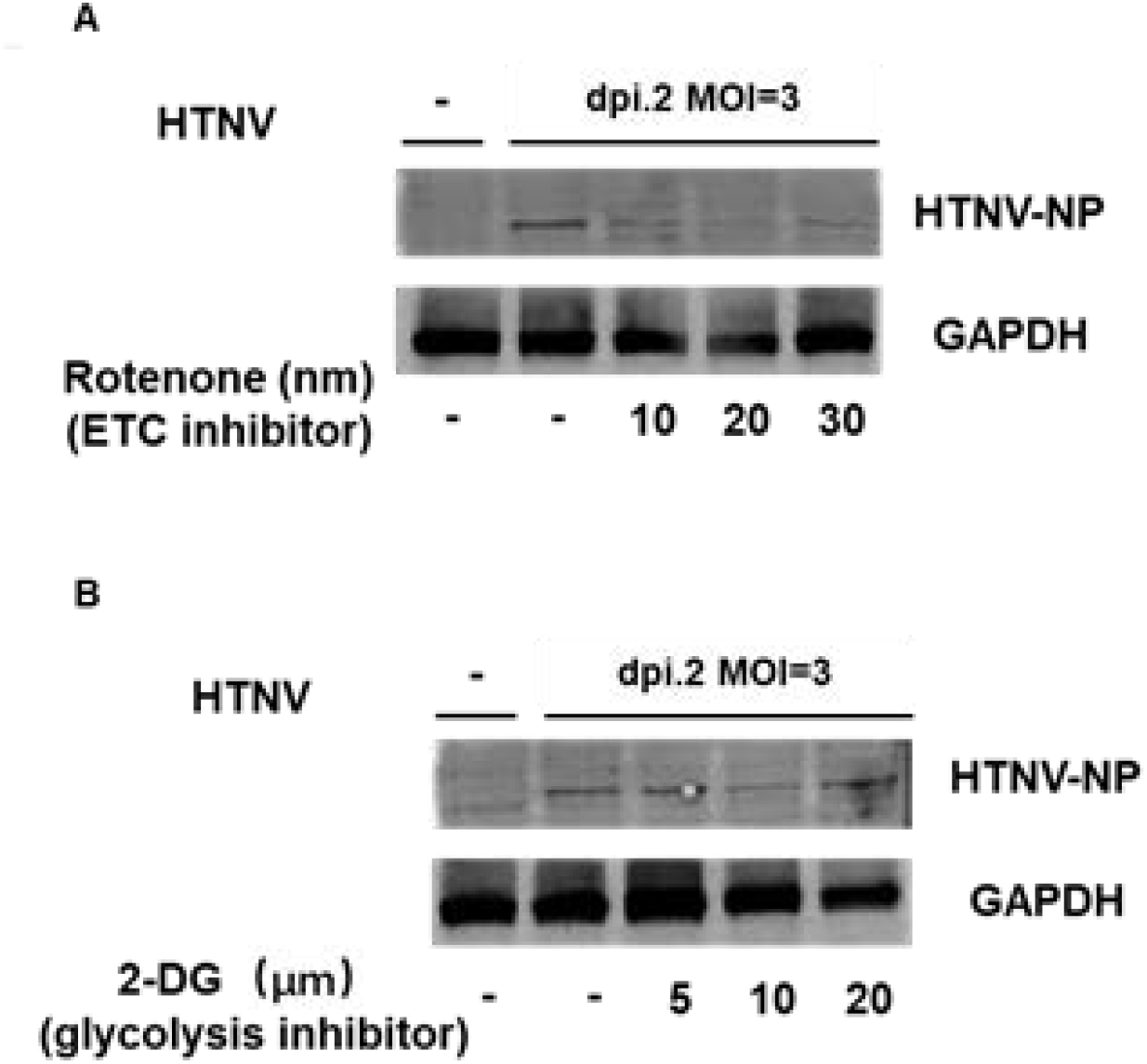
Rotenone can inhibit HTNV replication in A549 cells. We treated A549 cells with Rotenone (ETC inhibitor) or 2-DG (glycolysis inhibitor) before HTNV infection. Rotenone can inhibit HTNV replication effectively (A), but 2-DG rarely impair HTNV replication (B).

### HTNV infection induces mitochondrial biogenesis

Based on these results, we hypothesized that higher mitochondrial respiration activity after HTNV infection may accompanied with mitochondrial biogenesis enhancement. mtDNA transcription product includes 2 ribosomal RNAs, 22 tRNAs, and 13 mRNA which encoded essential subunits of mitochondrial respiratory complex^[29]^. Hence, we detected 13 mitochondrial mRNA transcription level after HTNV infection and found an increase of ND1, ND3, ND4, ND4L, ND5 (C I), CO III (C IV) and ATP6, ATP8(C V) (Figure 3A). BN-PAGE showed that mitochondria respiratory chain (RC) complexes I, IV, V expression were increased after HTNV infection (Figure 3B). Western blot also showed that mitochondria RC markers: NDUFB8 (C I), COI (C IV), UQCRC2(C III) and ATP5A (C V) expressions were enhanced obviously (Figure 3C). Within the most obvious enhancement of complex I and ATP synthase (complex V), there is no change of complex II expression level whose subunits were all encoded by nuclear gene. These results demonstrated mitochondrial biogenesis was increased after HTNV infection.

**Fig 3.**
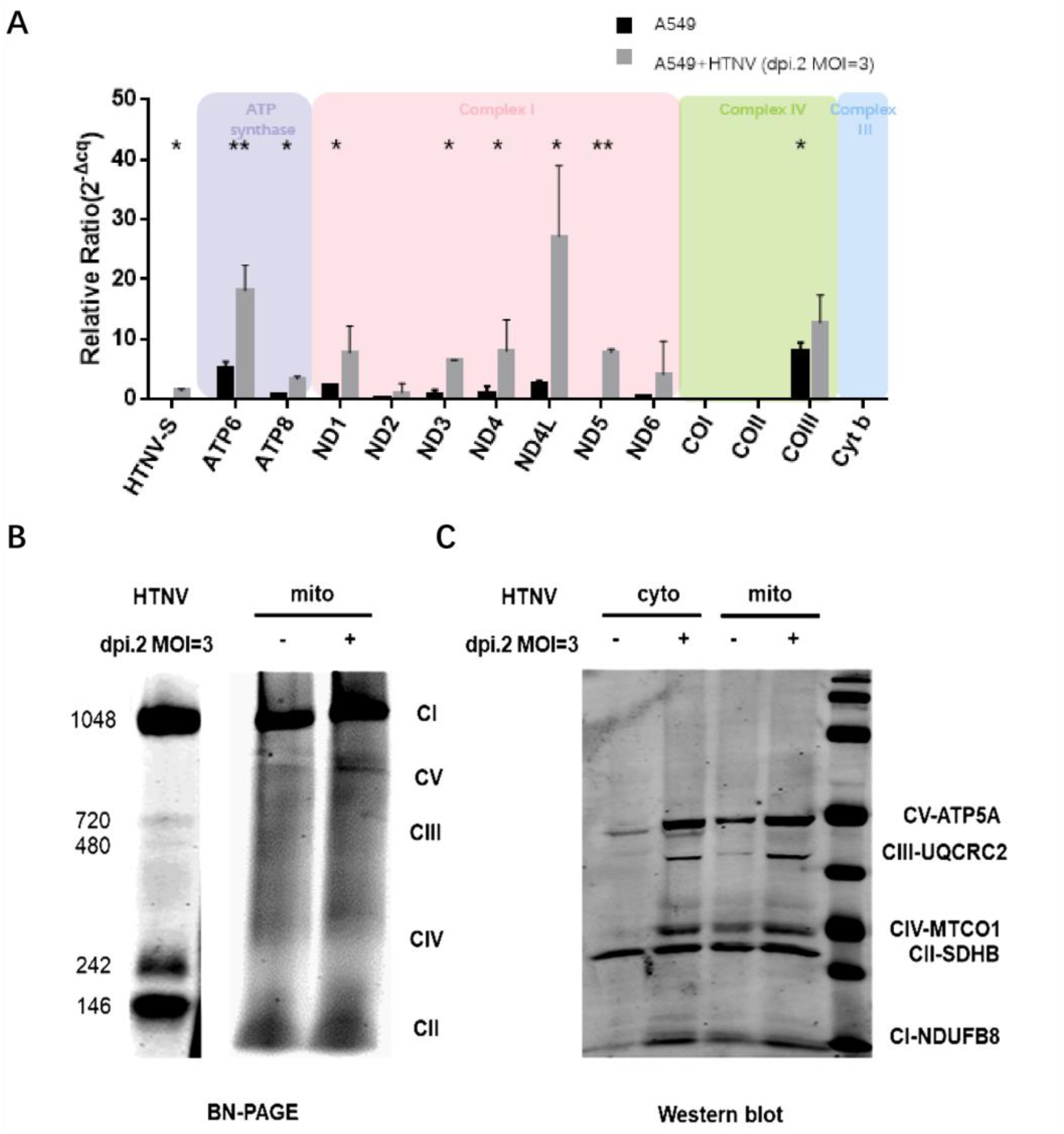
Mitochondrial biogenesis was increased after HTNV infection in A549 cells. (A) A549 cells were infected with HTNV at MOI of 3. 2 days after infection, the transcription level of mtDNA was measured by qRT-PCR. Values are means±SD (n= 3; *, P <0.01; **, P<0.001; ***, P<0.0001; Student’s t test, compared with the NC group). Mitochondria protein from A549 cells were isolated and detected by BN-PAGE (B) and western blot (C).

Besides, we observed mitochondria morphologic changes after HTNV infection. Using transmission electron microscopy (TEM), we found that after HTNV infection, mitochondria matrix became denser which means more biogenesis was happening at this situation (Fig 4).

**Fig 4.**
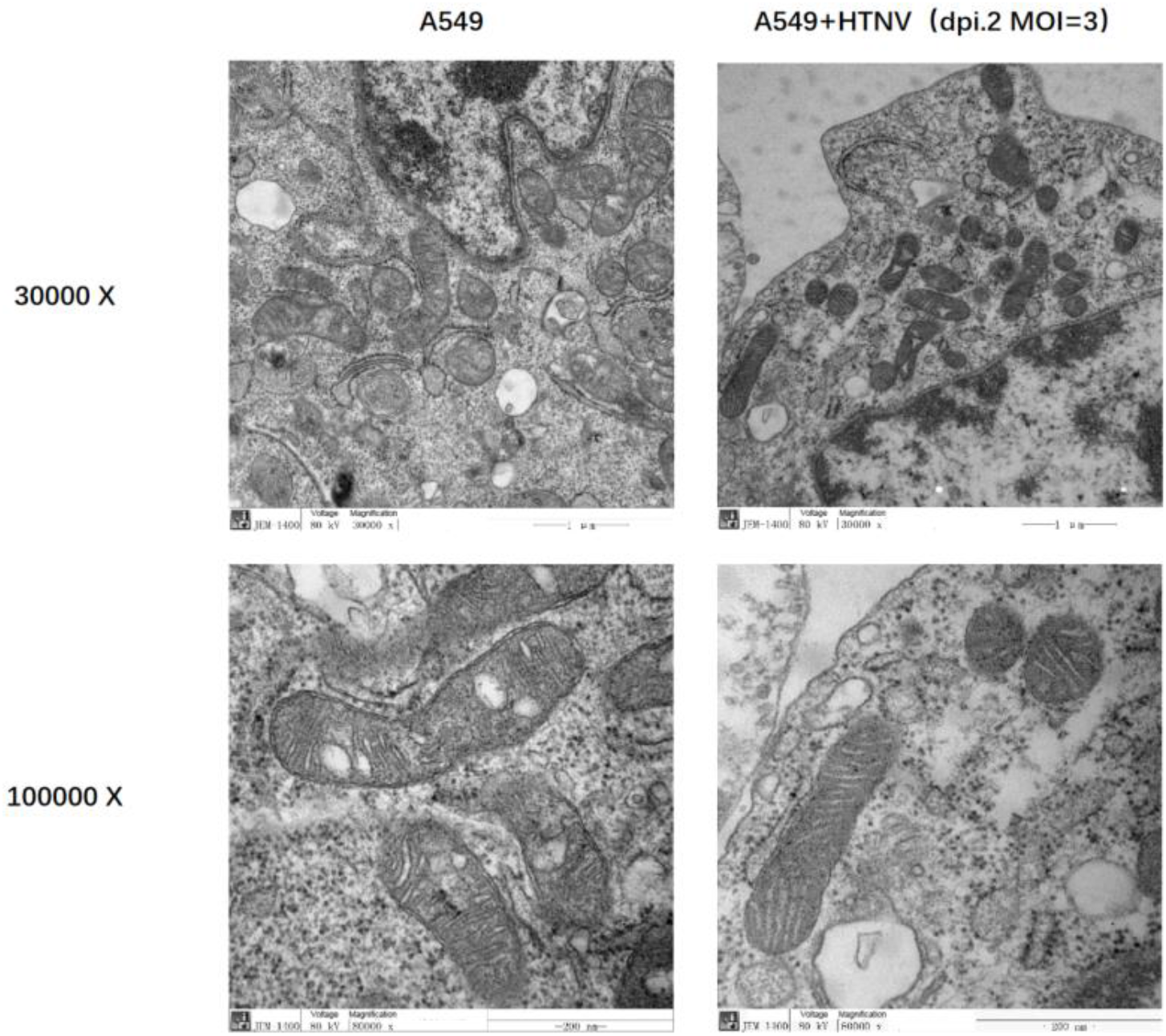
Mitochondria matrix density was increased after HTNV infection. 2 days after HTNV infection, cells were collected, fixed and sliced for TEM. Sections were observed using JEM-1220 transmission electron microscope (JEOL ltd, TOKYO, Japan). We can see a distinct density increase of mitochondria matrix after HTNV infection, and less vacuole which was cause by more crowded matrix after HTNV infection.

### AKT is activated and translocated to mitochondria after HTNV infection

To identify cellular signaling pathways that participating in the metabolic changes after HTNV infection, combined with our previous experiment results that HTNV infection did not inhibit cell proliferation but promoted cell proliferation slightly (Fig. S3), we focused on a key regulator of cell proliferation and metabolism, AKT. AKT is a major downstream effector of PI3K and can be phosphorylated at Thr^308^ and Ser^473 [30]^. During HTNV infection, AKT Thr^308^ and Ser^473^ were both phosphorylated at 24 hpi, and phosphorylation level of AKT ser^473^ was higher than AKT thr^308^ (Fig. 5A). Treating A549 cells with 40 nm BEZ (a potent inhibitor of PI3K) 4 hours before infection can completely inhibit AKT phosphorylation (Fig. 5A) and HTNV replication (Fig. 5A, B). These results showed that AKT activation was vital for HTNV replication.

**Fig 5.**
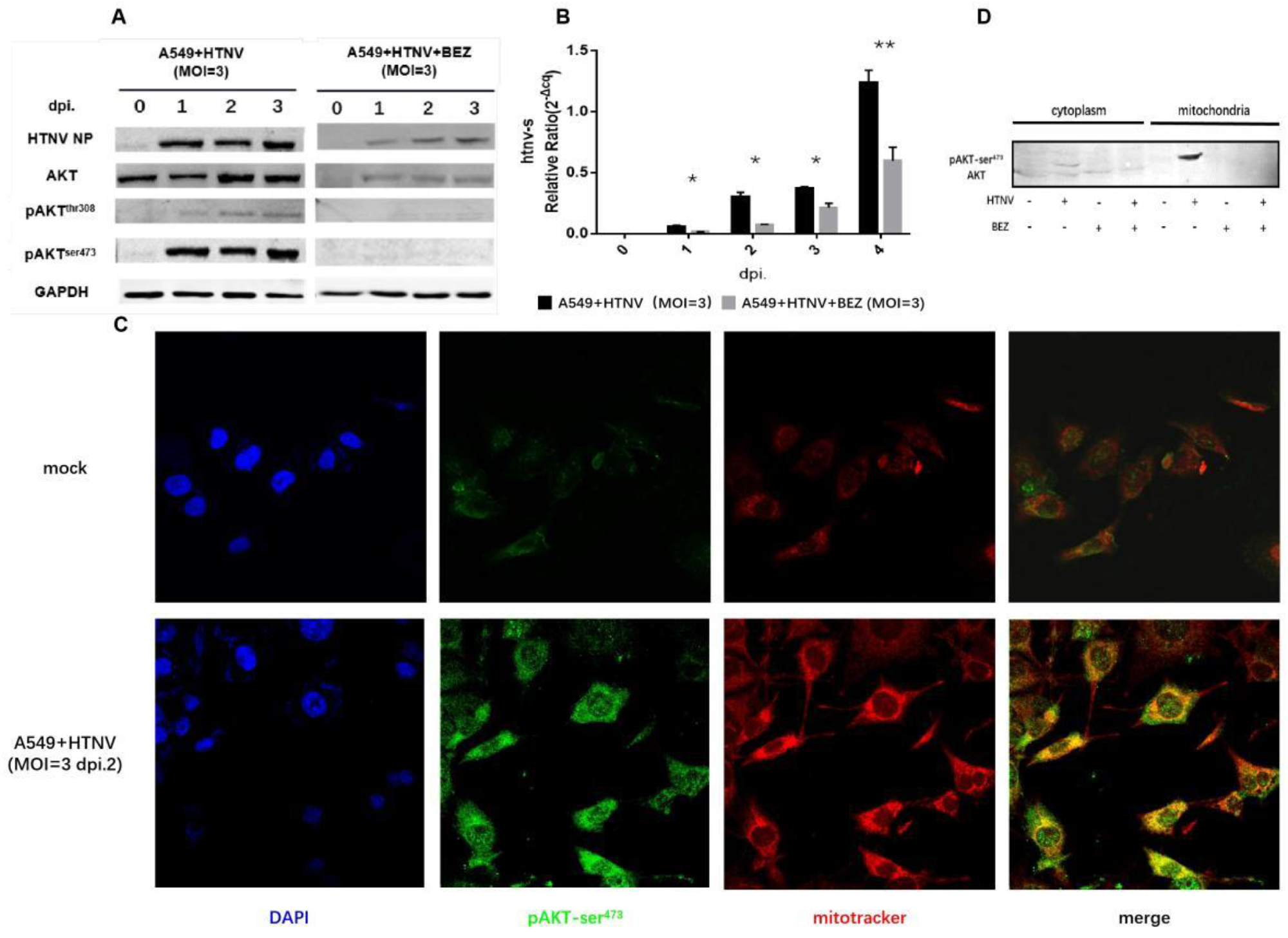
AKT activation and translocation after HTNV infection in A549 cells. A549 cells were infected with HTNV at MOI of 3 with DMSO or BEZ. Total protein was collected and tested via western blot (A) and HTNV-S segment transcription level was detected via qRT-PCR (B). AKT ser^473^ was phosphorylated quickly at early stage after HTNV infection. BEZ treatment in advance (4 hours before HTNV infection) can totally inhibit AKT phosphorylation and HTNV replication. As for pAKT-ser^473^ subcellular location after HTNV infection, Immunofluorescence assay (IFA) results showed that pAKT-ser^473^ (green) co-localized with Mitotracker (red) (C) and western blot showed an obvious pAKT-ser^473^ band in mitochondria fraction after HTNV infection.

As PH domain of AKT has high affinity with phosphate inositol, which allowed AKT translocate to any membrane structure, including mitochondria membrane^[23, 31]^. Based on this theory, we detect AKT subcellular location after HTNV infection. Confocal microscopy analysis showed appearance of pAKT-ser^473^ at mitochondria after HTNV infection (Fig. 5C). We then isolated mitochondria and cytoplasm and found that there existed pAKT-ser^473^ in mitochondria fractionation after HTNV infection (Fig. 5D), which was consistent with immunofluorescence results.

### Mitochondrial AKT modulated OXPHOS during HTNV infection

As pAKT-ser^473^ can translocate to mitochondria after HTNV infection, we want to know whether AKT play an important role on mitochondria function increase during HTNV infection. We found that mtDNA transcriptional level enhancement after HTNV infection did not appear if treating cells with BEZ in advance (Figure 6A), which suggested that if AKT activation was inhibited, HTNV infection cannot induce mitochondrial biogenesis enhancement.

**Fig 6.**
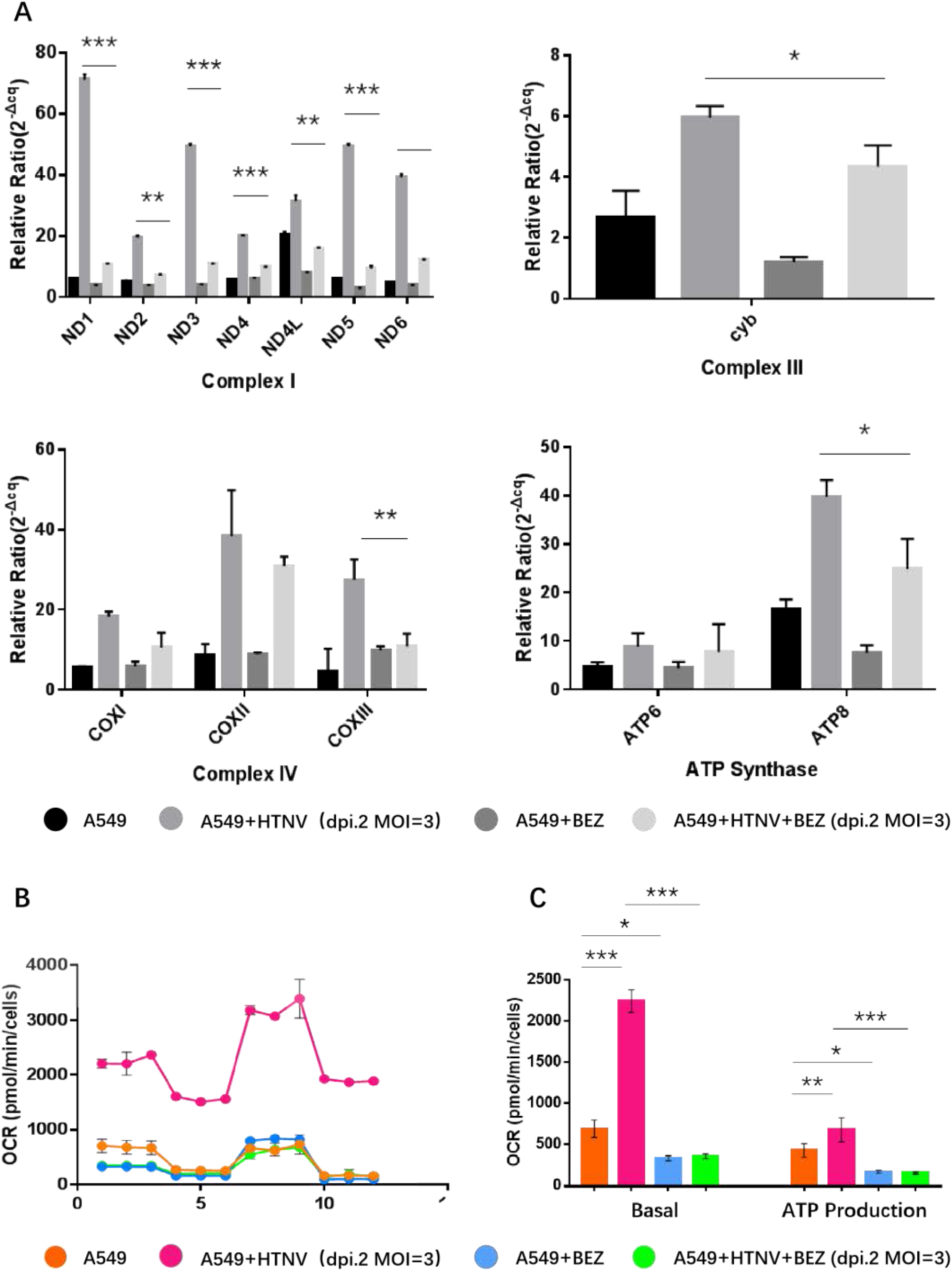
AKT activation promoted HTNV-induced mitochondrial function increase. A549 cells were treated with PI3K inhibitor BEZ (40 nm) 4 hours before infection, and then were infected with HTNV. The effect of BEZ on mtDNA transcription was detected after HTNV infection via qRT-PCR (A) (Values are means±SD (n= 3; *, P<0.01; **, P<0.001; ***, P <0.0001; Student’ s t test, compared with the NC group). Mitochondria respiratory capacity was also measured using the Seahorse XFe24 Mito Stress Test (B, C). Error bars indicated the standard errors of the mean (SEM) of at least triplicates. (*, P<0.01; **, P<0.001; ***, P<0.0001.)

Seahorse results showed that after HTNV infection, A549 cells without BEZ treatment, basal OCR (p < 0.0001) and ATP production (p < 0.001) were increased obviously (Figure 6B, C). But with BEZ treatment, even after HTNV infection, there is no statistically significant increase on basal OCR or ATP production level (Figure 6C), which meant that BEZ treatment can prevent the increase of basal OCR and ATP production caused by HTNV infection (p< 0.0001) (Figure 6C). Taken together, these results demonstrated that HTNV replication increase mitochondria OXPHOS via AKT activation, and this effect can be blocked by BEZ treatment.

### Phosphorylation of PNPT in mitochondria by AKT after HTNV infection

pAKT-ser^473^ accumulation in mitochondria after HTNV infection raised the question that if there existed any substrate that can be phosphorylated by AKT in mitochondria after HTNV infection. To address this issue, firstly, we analyzed mitochondrial proteins which have the consensus AKT substrate motif RXRXXT*/S*^[32]^ (where X represents any amino acid, and T* and S* represent phosphorylated threonine and serine respectively). After screening, 2 mitochondrial protein, PNPT (^12^RLRPLS^17, 26^RDRALT^31^) and NDUFB8 (^114^RNRVDT^119^) matched the AKT consensus phosphorylation motif.

Secondly, we used PAS antibody, which can recognize phosphorylated RXRXXT/S motif of AKT, to detect if this motif in NDUFB8 or PNPT was phosphorylated or not. We recruited PNPT protein from cytoplasm or mitochondria via anti-PNPT antibody and observed a strong PAS immunoreactivity of PNPT in HTNV-infected mitochondria pellet fraction (Figure 7A), which meant there existed a phosphorylation on RXRXXT motif of PNPT. There is no PAS reactivity of NDUFB8. We also found that the enriched pAKT-ser^473^ from HTNV-infected mitochondrial fraction can be recognized by anti-PNPT antibody (Figure 7B), which indicated an interaction between PNPT and pAKT-ser^473^ in mitochondria after HTNV infection, also implied that PNPT may be a potential substrate of AKT in mitochondria after HTNV infection.

**Fig 7.**
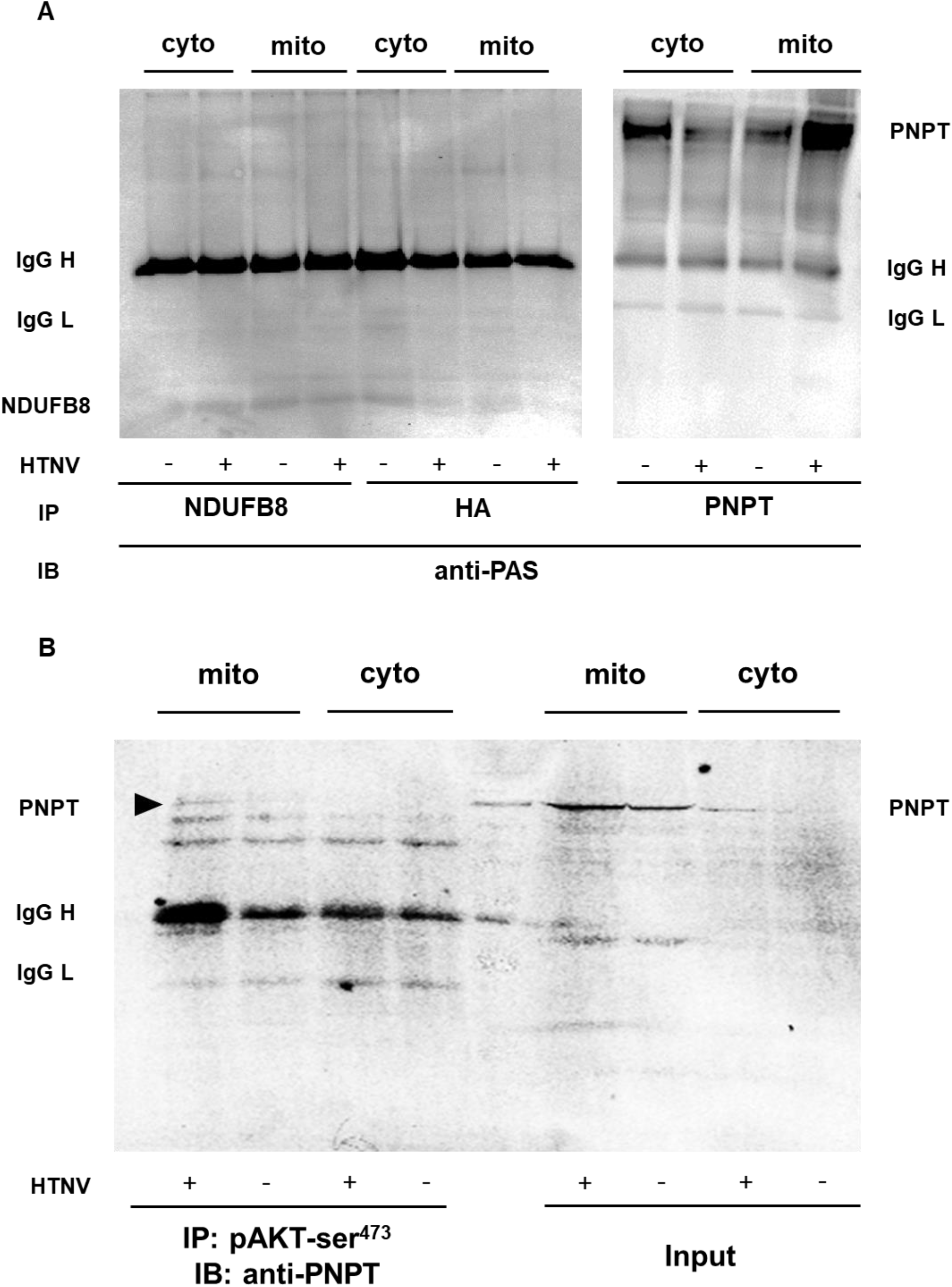
PNPT was a potential substrate of AKT in mitochondria after HTNV infection in A549 cells. A549 cells were infected with HTNV at MOI of 3. 2 days after infection, cytoplasm and mitochondria protein were isolated and co-incubated with anti-PNPT and anti-NUDFB8 antibody to enrich targeting proteins. Then the phosphorylation activity of RXRXXS/T were detected using PAS antibody. The results showed that after HTNV infection, RXRXXS/T motif of mitochondria PNPT was phosphorylated (A). Furthermore, we recruited pAKT-ser^473^ via pAKT-ser^473^ antibody and detected its interaction with PNPT. Results showed that PNPT had an interaction with pAKT-ser^473^ after HTNV infection in mitochondria (Arrow) (B).

## Discussion

Virus replication needs resources from host cells, how HTNV manipulate metabolic pathways of host cells to assist its own replication is still unclear. For the first time, we find that HTNV exploits mitochondria OXPHOS to assist its replication. Essentially, virus replication includes viral genome replication, transcription and viral protein expression, classical steps following Central dogma. Completion of these steps needs adequate biomaterial and ATP, which are all from host cells. As cell metabolism is a flexible process that adapts to various stimuli, such as infection, substrate supply, hormonal and growth factor stimulation, thus, we choose 3 different cells to comprehensively evaluated their bioenergetic profiles after HTNV infection. Via seahorse XFe24 Bioanalyzer, we found that after HTNV infection, mitochondria OXPHOS level and ATP production were increased, meanwhile, inhibiting mitochondria OXPHOS by rotentone treatment can inhibit HTNV replication. Instead, using glucose uptake inhibitor 2-DG had no influence on HTNV replication. These results showed that HTNV replication relied on mitochondria OXPHOS from host cells. And then we verified that mitochondrial biogenesis was enhanced after HTNV infection.

S. Karniely et.al^[7]^ raised a view that viruses which can cause CPE after infection prefer glycolysis pathway, while those viruses do not cause CPE may choose OXPHOS to ensure ongoing supply of energy during its long replication cycle: They found that although HSV and HCMV all belong to Herpesviridae, HSV can survive in host cells for only 1-3 days and can cause severe cytopathic effect and HSV encoded non-structural protein US3 can inhibit mitochondria OXPHOS level^[33]^; HCMV can proliferate in host cells for 7-10 days, increase cellular mitochondria biogenesis and function^[33]^. Our findings verified this theory but how HTNV mediated cellular metabolism during its replication was still unknown.

Some viruses can promote AKT activation to inhibit cell apoptosis to maintain virus replication in host cells. For example, HBx protein encoded by HBV can activate AKT and phosphorylate Hepatocyte Nuclear Factor 4 (HNF4) to inhibit cell apoptosis caused by HBV infection, maintain HBV chronic infection status, but does not induce cytopathic effect in host cells^[35]^. Influenza virus^[36]^ and Respiratory Syncytial Virus^[37]^ can also activate AKT signal pathway to inhibit cell apoptosis.

Considering that HTNV replication can promote cell proliferation slightly, we wondered if AKT was involved in th<u>e</u> energy utilization process during HTNV infection and we found that after HTNV infection, AKT was activated and inhibition of AKT activation by BEZ can obviously inhibit HTNV replication.

Although AKT is mostly reported to mediate glucose metabolism, there are also many studies showed that AKT can enhance mitochondria function in some particular conditions^[25, 38-40]^. We found that after HTNV infection, AKT can be phosphorylated and pAKT-ser^473^ can translocate to mitochondria. We also found that PI3K-AKT inhibitor BEZ235 can inhibit AKT activation and translocation to mitochondria and decrease mitochondria OXPHOS level after HTNV infection.

The finding that pAKT-ser^473^ can translocate to mitochondria after HTNV infection was so interesting that we thought that AKT can phosphorylate some mitochondrial protein to mediate mitochondria OXPHOS during this process. We found that after HTNV infection, pAKT-ser^473^ can interact with PNPT, a mitochondria protein which mediate mtRNAs turnovers and its dysfunction can leads to mitochondrial respiratory function decrease: PNPT knockout in mouse embryonic fibroblasts resulted in the loss of mtDNA and perturbation in mtDNA transcription^[41]^; Patients with PNPT missense mutation was born with severe encephalomyopathy, choreoathetotic movements and respiratory-chain defects^[42]^. PNPT mutations genetically link to hereditary hearing loss, encephalomyopathy, and axonal and auditory neuropathy, gut disturbances, chorioretinal defects, Leigh syndrome, and delayed myelination ^[43-50]^. Thus, based on our results, we can speculate that after HTNV infection, AKT was activated and translocated to mitochondria, where it phosphorylated PNPT, and whose phosphorylation may promote mitochondria respiratory function.]

In conclusion, it is the first time to find a specific phenomenon that HTNV infection utilize mitochondria OXPHOS in host cells to assist its replication, and its underlying mechanism could be that HTNV infection can activate AKT-PNPT signal pathway in mitochondria to enhance mitochondria biogenesis and respiration function. It is also the first time to find a new substrate of AKT in mitochondria. Our findings provide a new thought for anti-viral drug development, it is also a meaningful complement for HTNV biological characteristics.

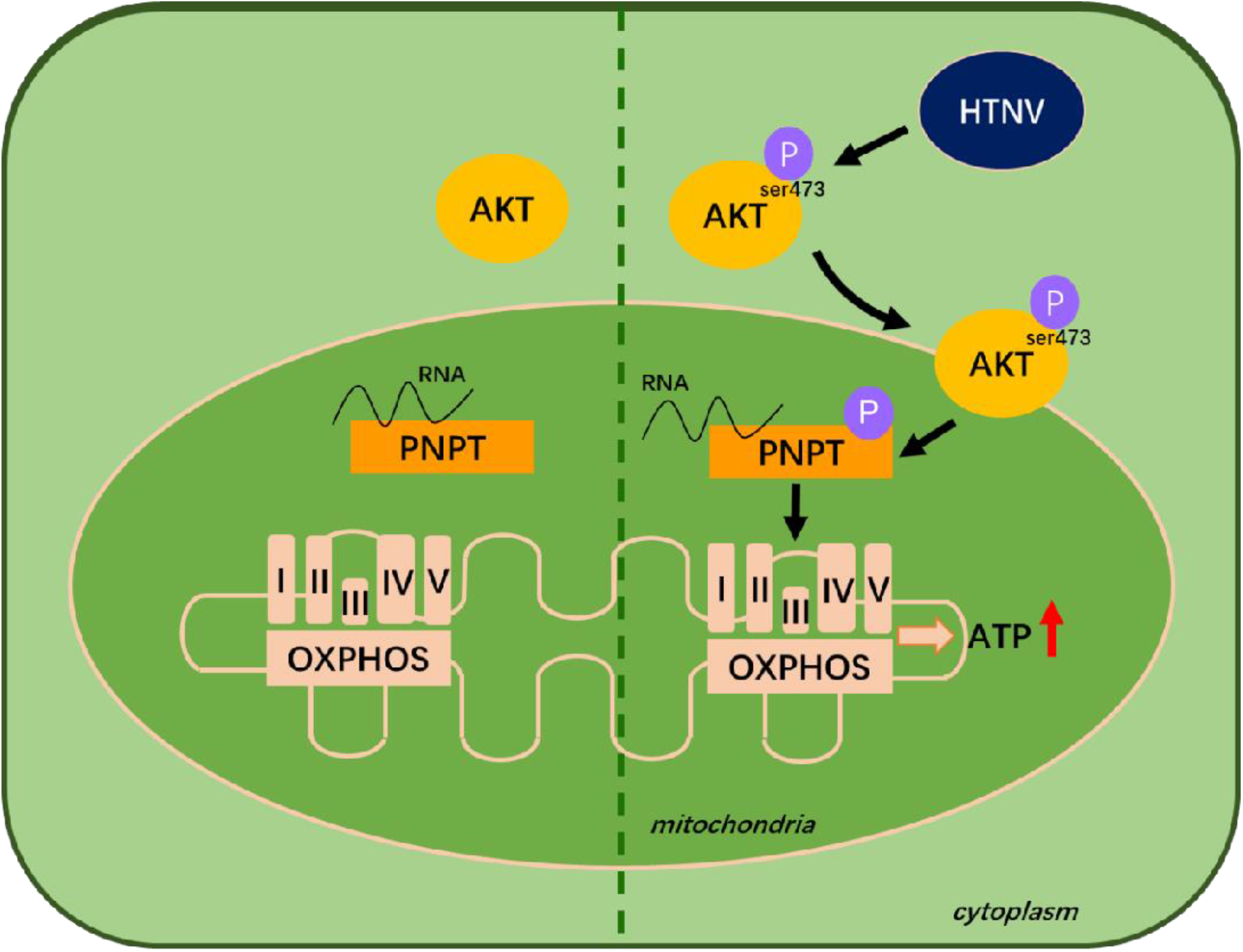

## ACKNOWLEDGMENTS

This work was supported by the Foundation for scientific and technological project in Shaanxi Province (No.2019ZDLSF02-03,No.2021JM-219) and the funding of Air Force Military Medical University (No.2018JSTS08).

---

**Fig S1.**
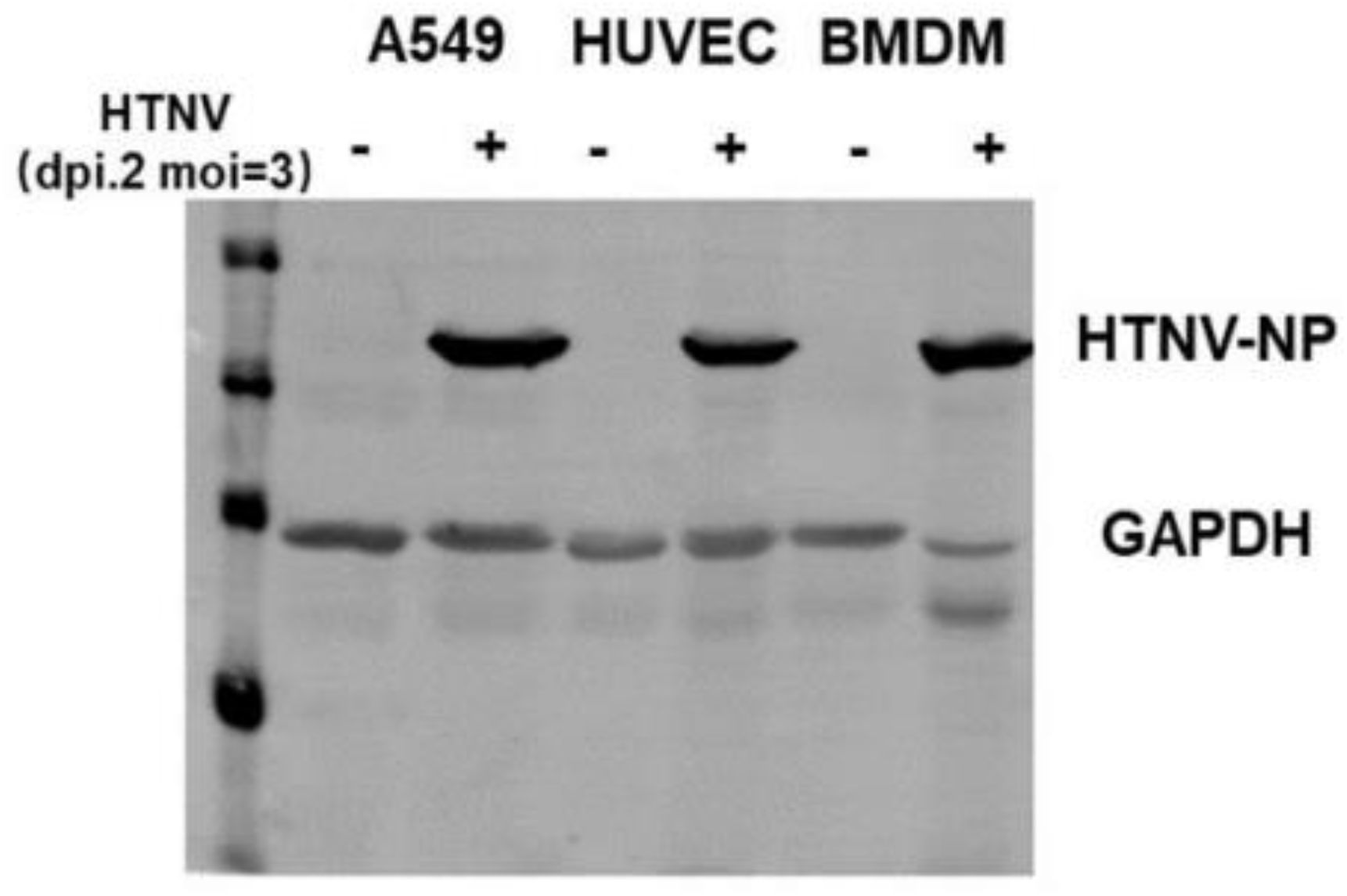
HTNV-NP expression after infection in different cells. A549, HUVEC or mouse BMDM cells were infected with HTNV at MOI=3, 2 days after infection viral nucleocapsid protein was detected by western blot with anti HTNV-NP mAb 1A8.

**Fig S2.**
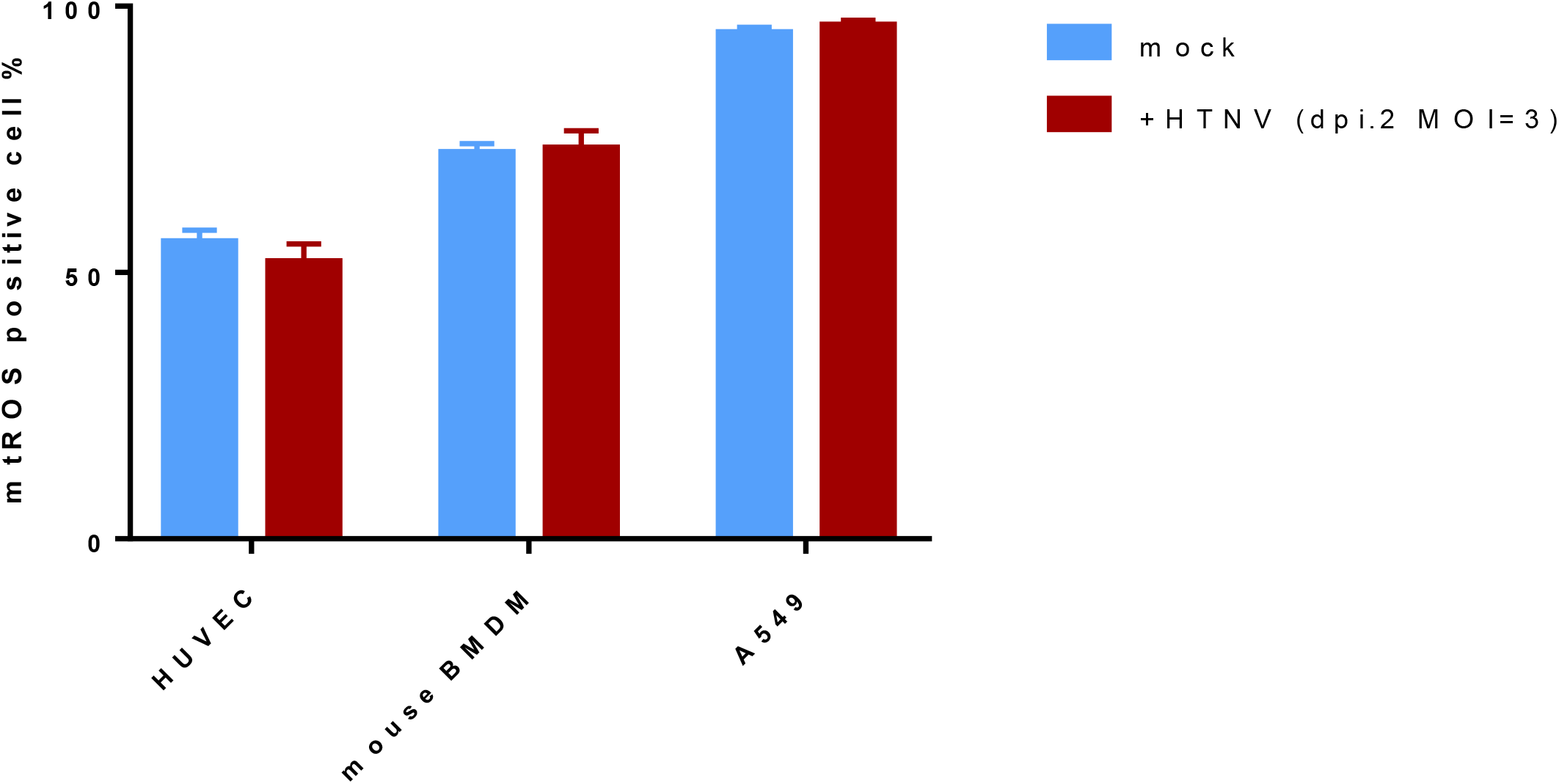
Mitochondiral ROS did not change after HTNV infection. Mitochondrial ROS was detected by Flow Cytometry with the probe MitoTracker red CMXRos (ThermoFisher Scientific Technologies). Cells were incubated with MitoSox (2.5mM) for 15 min and detected by BD caliber flow cytometer. The results were analyzed by FlowJo v9.3.2and there is no statistical difference of mtROS in mock or HTNV-infected A549 cells. (Student’s t test, n=3, compared with the mock group).

**Fig S3.**
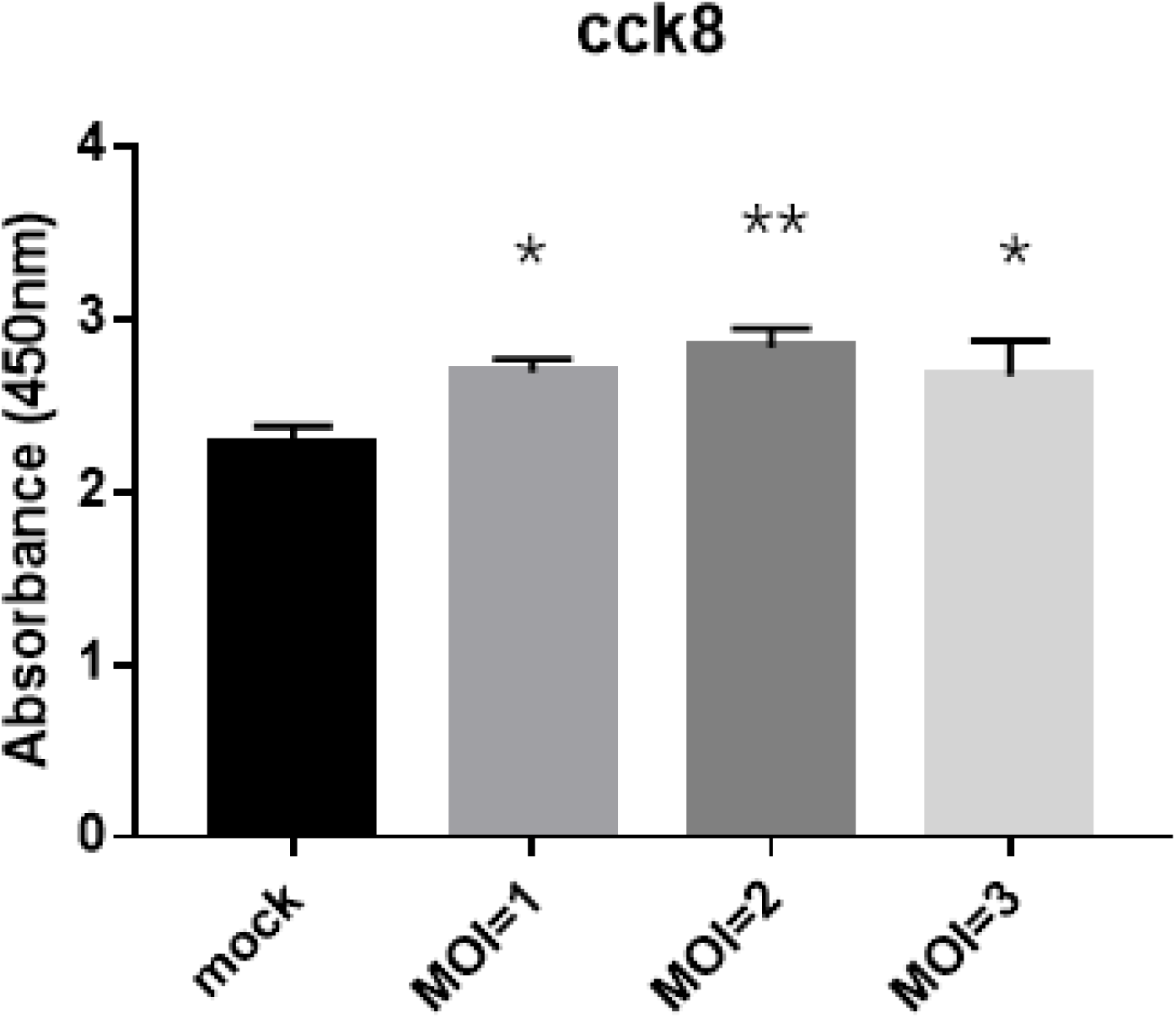
HTNV infection promotes cell proliferation. A549 cells were infected with HTNV at MOI of 1, 2, 3 respectively and detected by CCK8 assay. Briefly, after 2 days infection of HTNV, 10 μl CCK8 was added and incubated for 4 hours and the absorbance was measured using a Microplate Reader (Bio-Rad) at wavelength of 450 nm. (Student’s t test, n=3, compared with the mock group).

## Materials and Methods

### Cell Culture

Vero-E6 and A549 cells were purchased from American Type Culture Collection (ATCC), and were maintained in Dulbecco’s modified Eagle’s medium (DMEM) supplemented with 10% FBS, 1% penicillin and 100μg/mL streptomycin. Human Umbilical Vein Endothelial Cells (HUVECs) were purchased from ScienceCell and were maintained in Endothelial Cell Medium (ECM) supplemented with 10% FBS, 1% penicillin and 100μg/mL streptomycin. Primary mouse bone marrow derived macrophages (BMDMs) were isolated from C57 mouse thigh bones, maintained in DMEM supplemented with 10% FBS and 10ng/mL Macrophage-Colony Stimulating Factor (M-CSF). All cells were grown at 37°C and 5% CO_2_ in a humidified incubator.

### HTNV strain and virus infection

HTNV strain 76-118 was isolated from the brain by intracranial inoculation of Kunming Mouse (KM) litters and followed by propagating in vero E6 cells. After propagation in vero E6 cells, HTNV was titrated by foci forming unit (FFU) assays in Vero E6 cell monolayers.

### FFU/mL=average Foci counts in each cell × dilution/virus volume added in each well

For infection, cells with a confluence of 70 to 80% in 6, 24, or 96 wells or 35-mm dishes were rinsed twice with Dulbecco’s phosphate-buffered saline (DPBS; HyClone), followed by the addition of HTNV that had been diluted to the desired multiplicity of infection (MOI). In our experiments, all cells were either mock infected (control) with medium excluding virus or infected with HTNV 76-118 strain at a MOI of 3.

### MOI=FFU/cell counts

After incubation for 90 min at 37°C, the supernatant was discarded and maintained in medium supplemented with 2% FBS.

### Measurements of mitochondria oxidative phosphorylation (oxygen consumption)

Mammalian cells get ATP from mitochondria OXPHOS and cytoplasm glycolysis. Seahorse extracellular flux analyzer can be used to measure oxygen consumption rates (OCR) and Extracellular Acidification Rate (ECAR) in cells.

For bioenergetic flux analyses, approximately 15,000 A549 cells were used to seed the wells of XF cell culture microplates (Seahorse Bioscience, US). On the analysis day, the incubation medium for each well was aspirated, the adherent cells were washed, and incubated in serum-free, pyruvate-free, buffer-free DMEM containing 25 mM glucose. After the switch to unbuffered medium, the microplates were placed in a 37°C, non-CO_2_ incubator for 1 h and then transferred to the microplate stage of a Seahorse XF24 flux analyzer (Seahorse).

Inhibitors during OCR measurement were used at the following concentrations: Oligomycin (1.0 µM), Carbonyl cyanide 4-trifluoromethoxy-phenylhydrazone (FCCP) (0.75 µM), Antimycin A (1.5 µM) and Rotenone (2 µM).

To account for variations in cell number brought about by drug-induced effects on proliferation or cell death, we used crystal violet for cell counting and data standardization, then analyzing OCR data via Wave software.

### Mitochondrial isolation

The isolation procedure was performed using Minute^™^ mitochondria isolation kit for cultured cells, according to the manufacturer’ s protocol. The whole procedure was performed on ice bath in order to avoid protein inactivation or degradation. For further protein phosphorylation measurement, all the isolation reagents were added Protease and Phosphatase Inhibitors (Thermo) in advance.

Briefly, cells were digested by trypsinization, harvested, and centrifuged at 800 g for 5 min. 100,000,000 cells were collected for each mitochondrial isolation procedure. Cells were washed twice in ice-cold TBS before beginning isolation procedure. Cells were homogenized on ice in 500 μL buffer A. Incubating the cell suspension on ice for 30 min and vortex vigorously for 20-30 seconds every 10 min. Then capping the filter cartridge and centrifuge at 16,000 g for 30 seconds. Discard the filter and resuspend the pellet by vortexing briefly. Centrifuge at 700 g for 1 min, carefully transfer the supernatant to fresh 1.5 ml tube and add 300 µl buffer B to the tube (the supernatant to buffer B ratio is 1:1.2). Mix by vortexing for 10 seconds. Centrifuge at 16,000 g for 30 min. Remove the supernatant completely and resuspend the pellet in 200 µl buffer B by repeat pipetting up and down followed by vigorously vortexing for 10-20 seconds. Centrifuge the tube at 8,000 g for 5 min. Transfer the supernatant to a fresh 2.0 ml tube; add 1.6 ml cold PBS to the tube and centrifuge at 16,000 g for 30 min. Discard the supernatant and save the pellet (isolated mitochondria).

### BN-PAGE

Lysing cells with Native lysis Buffer (Sangon): 0.1 g fresh mitochondria protein added with 1 mL Solution A, 1 μL Solution B and 10 μL Solution C,vortexing violently and dispersing completely, lysing on ice for 20 min. Then centrifuged at 16,000 g for 25 min, extracted supernatant and determined protein concentration using a bicinchoninic acid (BCA) assay kit (Thermo, USA). Mix 50 μg protein with 2× NativePAGE sample buffer (Sangon).

Load sample into the gel, then fill the inner chamber with about 180 ml of dark blue cathode buffer, and then fill the outside chamber with about 600 ml of running buffer. Turn on the power supply and run the gel at 150 V for 30 min, 250 V for 120 min. After electrophoresis, taking the gel out of the cassette and wash it with water. Then incubate the gel in transfer buffer for 15 min with gentle shaking.

### Assessment of mitochondrial respiratory chain protein

#### (1) Protein sample preparation

Cells were washed twice with ice-cold TBS and lysed with RIPA buffer containing protease inhibitor and phosphatase inhibitor (Thermo, US). Protein concentration was determined using a BCA assay kit (Thermo, USA). 30 mg protein was then boiled at 95°C for 10 min.

#### (2) SDS-PAGE and Western blot analysis

The lysates were separated by 10% SDS-PAGE and transferred to polyvinylidene fluoride (PVDF) membranes (Millipore). The membranes were incubated with the primary antibodies, followed by secondary antibodies labeled with infrared dyes (Li-Cor Biosciences, Lincoln, NE, USA). The signals on the PVDF membrane were visualized using an Odyssey infrared imaging system (Li-Cor Biosciences, Lincoln, NE, USA).

### RNA extraction and quantitative real-time RT-PCR analysis

Total RNA was purified from cells using a Tiangen Easy Fast RNA Extraction kit and reverse-transcribed using Transcriptor First Strand cDNA Synthesis kit (Roche). Expression of mtDNA encoded genes were determined by quantitative real-time RT-PCR using SYBR Green PCR Master Mix (Roche Applied Science). The primers were listed as followed.

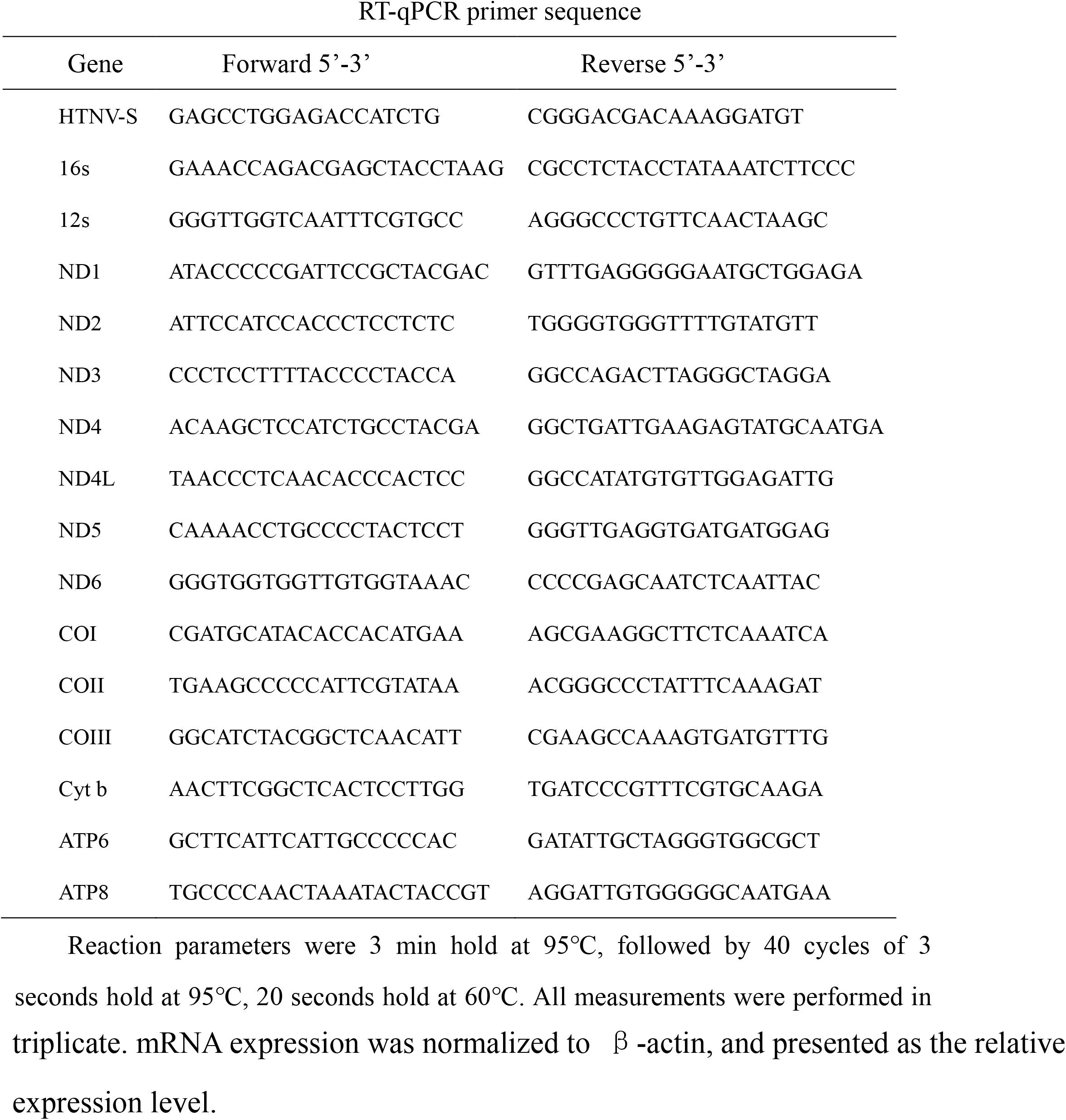

### immunofluorescence assays (IFA)

A549 Cells grew on coverslips in 24-well cell culture plates. 2 days after infection, cells were washed with Tris HCl buffered saline (TBS) and fixed in 4% formaldehyde (Sangon) for 20 min at room temperature. After washing 3 times with TBS, cells were permeabilized with 0.1% Triton X-100 (Sangon) in TBS for 20 min at room temperature and then blocked with TBS containing 3% BSA (Sangon) at room temperature for 1 h. Cells were then incubated with Phospho-Akt (Ser473) (D9E) XP® Rabbit mAb (1:1000) (Cell Signaling Technology)overnight at 4°C, washed 3 times with TBS added with :0.1% Triton (TBST), and incubated with Fluorescein (FITC)/Cy3-conjugated Goat Anti-Rabbit IgG(H+L)(Proteintech Group, Inc). Cell nuclei were stained with DAPI (4=,6-diamidino-2-phenylindole) (1:10000) and incubated for 10 min in dark place. After that, washing 5 times with TBST. Finally, the samples were observed using a confocal fluorescence microscope FV1000 (Olympus).

### Coimmunoprecipitation (Co-IP)

A549 cells were mock infected or infected with HTNV at MOI of 3, mitochondria and cytosol were isolated and lysed with IP lysis buffer (with protease inhibitor and Phos-STOP Phosphatase Inhibitor Cocktail) at 2 dpi. The lysates (200 μg/mL) were incubated with PNPT1 Polyclonal antibody or NDUFB8 Polyclonal antibody (Proteintech Group, Inc) or pAKT-ser^473^ polyclonal antibody (1:100) respectively and incubation buffer at room temperature for 2 h in spin columns. Meanwhile, control IgG was added to the lysate as a negative control. Protein A/G Magnetic beads (MCE) were then added, incubated at 4°C overnight, and then eluted with elution buffer. After centrifugation at 1,000 rpm at 4°C for 10 min, magnetic beads were resuspended in 30 μL 4× sample buffer, and boiled for 5 min. Then the samples were detected via western blot by PAS antibody (Proteintech Group, Inc) (1:1000).

### Transmission Electron Microscope (TEM)

At 2 dpi, mock and HTNV-infected A549 cells were collected and fixed with 2.5% glutaraldehyde in phosphate buffer at 4°C overnight. Cells were then embedded in Epoxy resin and sliced for electron microscopy. Sections were observed on a JEOL 1200 electron microscope (JEOL Ltd., Tokyo, Japan) operated at 80KV.

### Reagents

Abs against GAPDH, PNPT, HRP-conjugated goat anti-mouse IgG (H+L) and FITC-conjugated goat anti-mouse IgG (H+L) were purchased from Proteintech Group, Inc; anti Phospho-Akt Substrate (RXXS*/T*) (PAS), Akt (pan) (11E7) Rabbit mAb, Phospho-Akt (Ser473) (D9E) XP® Rabbit mAb, Phospho-Akt (Thr308) (D25E6) XP® Rabbit mAb, Phospho-Akt Substrate (RXXS*/T*) (110B7E) Rabbit mAb (HRP Conjugate) and Phospho-Akt Substrate (RXXS*/T*) (110B7E) Rabbit mAb (Sepharose®Bead Conjugate) were purchased from Cell Signaling Technology; Total human OXPHOS Antibody Cocktail was purchased from Abcam; IRDye® 680RD Goat anti-Rabbit IgG (H + L) and IRDye® 680RD Goat anti-Mouse IgG (H + L) were purchased from LI-COR Biosciences; MitoTracker™ Red FM was purchased from ThermoFisher Scientific Technologies. Rotenone and 2-DG were purchased from MCE and were dissolved in DMSO at a stock concentration. BEZ235 was purchased from Selleck and was dissolved in DMF at a stock concentration of 1 mM. BD ELISpot Chromogen Solution was purchased from BD Biosciences.

### Statistical analysis

OCR data was analyzed via Wave software. For all other experiments, data were analyzed in Prism7 using tests as indicated in the figure legends.

